# Breast cancer-associated skeletal muscle mitochondrial dysfunction and lipid accumulation is reversed by PPARG

**DOI:** 10.1101/2020.04.05.026617

**Authors:** Hannah E. Wilson, David A. Stanton, Emidio E. Pistilli

## Abstract

The peroxisome-proliferator activated receptors (PPARs) have been previously implicated in the pathophysiology of skeletal muscle dysfunction in women with breast cancer (BC) and in animal models of BC. Here, we sought to describe the metabolic alterations induced in skeletal muscle by BC-derived factors in an *in vitro* conditioned media (CM) system and hypothesized that BC cells secrete a factor that represses PPAR-gamma (PPARG) expression and its transcriptional activity, leading to downregulation of PPARG target genes involved in mitochondrial function and other metabolic pathways. We found that BC-derived factors repress PPAR-mediated transcriptional activity without altering protein expression of PPARG. Further, we show that BC-derived factors induce significant alterations in skeletal muscle mitochondrial function and lipid metabolism, which are rescued with exogenous expression of PPARG. The PPARG agonist drug rosiglitazone was able to rescue BC-induced lipid accumulation, but did not rescue effects of BC-derived factors on PPAR-mediated transcription or mitochondrial function. These data suggest that BC-derived factors induce deficits in lipid metabolism and mitochondrial function via different mechanisms that are both related to PPARG signaling, with mitochondrial dysfunction likely being altered via repression of PPAR-mediated transcription, and lipid accumulation being altered via transcription-independent functions of PPARG.

## INTRODUCTION

Breast cancer (BC)-associated skeletal muscle fatigue is a chronic problem among BC survivors, being reported by a majority of patients both prior to and after receiving anti-cancer therapies (1-5). Recent studies show that deficits in muscle function predict shorter survival in cancer, perhaps due to the fact that fatigue is known to reduce a patient’s tolerance to anti-cancer therapies (3,6-10). Therefore, improving muscle function in BC patients has the potential to improve both quality of life and survival in the most commonly diagnosed cancer type in women.

The peroxisome-proliferator activated receptors (PPARs) are lipid sensing, ligand activated transcription factors previously implicated in a variety of pathologies, including cancer, atherosclerosis, Alzheimer’s disease, type 2 diabetes mellitus, and others (11-14). These transcription factors aid in regulating whole body energy homeostasis via regulation of genes involved in lipid metabolism and mitochondrial functions (11,15-19). Recently, downregulation of PPAR-gamma’s (PPARG) transcriptional activity in skeletal muscle has been identified as a potential central regulator of the increased muscle fatigue experienced by women with BC and recapitulated in the patient-derived orthotopic xenograft model of BC (20). However, the mechanism by which PPARG downregulation induces muscle dysfunction has not yet been explored. We hypothesized that BC cells secrete a factor that represses PPARG expression and its transcriptional activity, leading to downregulation of PPARG target genes involved in mitochondrial function and other metabolic pathways.

## RESULTS

### BC-CM represses PPARG expression and PPAR-mediated transcription

Because PPARG is downregulated in the skeletal muscle of women with BC and in mouse models of BC (20), we first investigated the capacity of BC-derived factors to downregulate *Pparg* mRNA expression in an *in vitro* model system. To test this, we collected CM from BC or control cell lines and applied this CM to differentiated skeletal muscle cells in culture (**Figure 1A**). Cells and large debris were removed from the media via gentle centrifugation and decanting, and the conditioned media was diluted in 1:3 in fresh growth media to correct pH and overall nutrient content. Importantly, all CM was applied to myotubes within 2 hours of collection to ensure that biologically active components would not degrade or be damaged by freezing prior to application. Using this model system, we found that CM from BC cell lines significantly downregulated *Pparg* mRNA expression (**Figure 1B**) and several of its purported transcriptional targets (21) in differentiated skeletal muscle after as little as two hours of CM exposure, compared to media conditioned by skeletal muscle myoblasts (**Figure 1C**). However, no significant change in PPARG protein expression was detected (**Figure 1D**), suggesting that the downregulation of PPARG target genes’ mRNA is due to a repression of PPAR-mediated transcriptional activity, rather than a down-regulation of PPARG protein abundance. To directly address this discrepancy, we generated a reporter cell line stably expressing a PPAR-responsive reporter construct driving expression of GFP. Using this system, we have consistently observed a significant reduction in GFP intensity in response to BC-CM, compared to CM from the reporter cells themselves. In contrast, CM from a non-tumorigenic mammary epithelial cell line does not repress GFP expression (**Figure 1E**).

**Figure 1:**
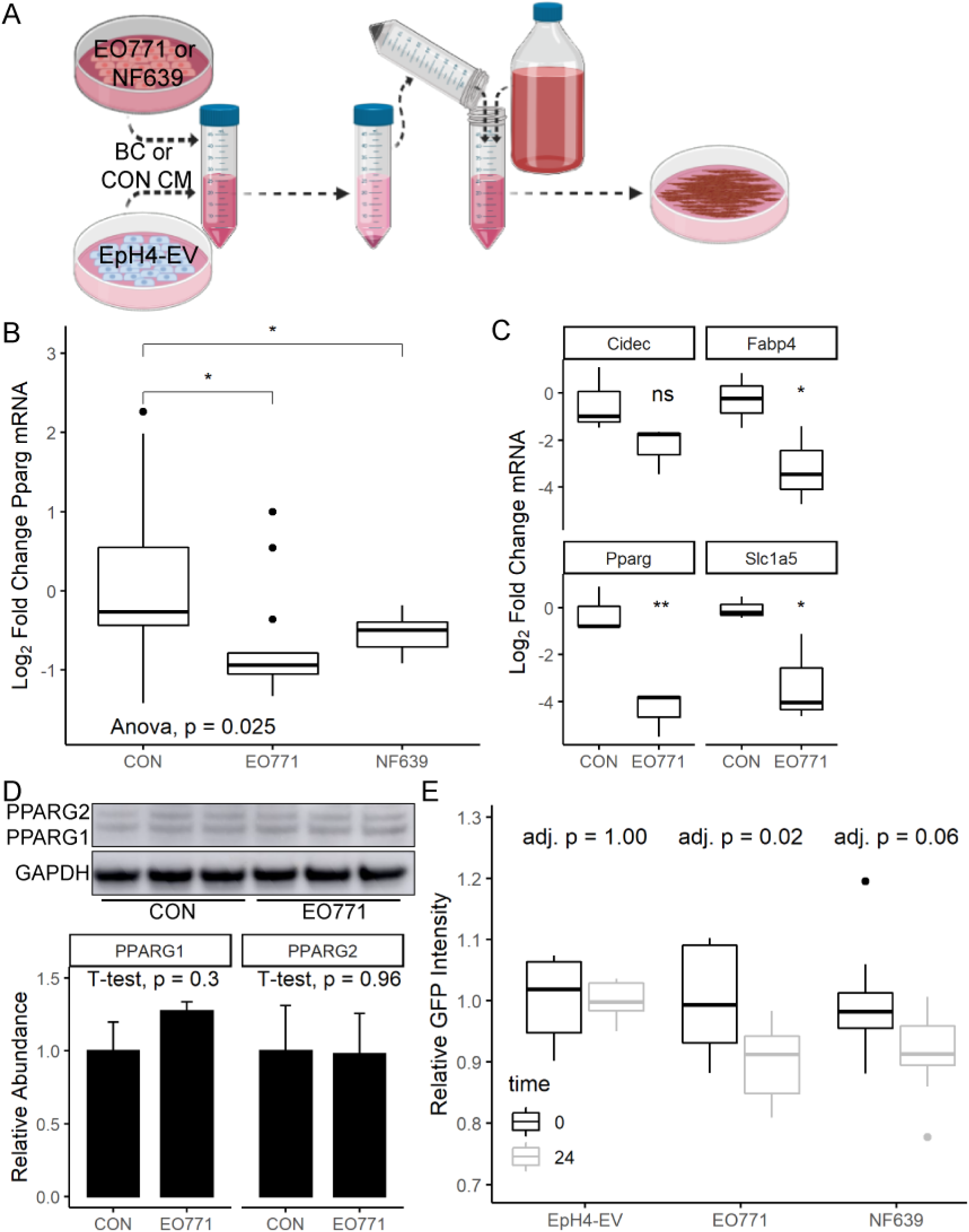
BC-CM represses PPARG expression and PPAR-mediated transcription. 48-hour conditioned media was collected from BC (EO771 or NF639) or control cell lines (EpH4-EV, C2C12 or HEK293), centrifuged to pellet large debris, decanted, and diluted in fresh complete growth media prior to application on recipient cells. Figure created using BioRender (**A**). *Pparg* mRNA expression in 5 day differentiated C2C12s exposed to 48-hour CM for 24 hours, quantified using qRT-PCR, n_CON_ = 12, n_EO771_ = 6, n_NF639_ = 5 (**B**). mRNA expression of PPARG target genes in 5 day differentiated C2C12s exposed to 48-hour CM for 2 hours, quantified using qRT-PCR, n_CON_ = 3, n_EO771_ = 3 (**C**). Western blotting analysis of PPARG protein expression in 5 day differentiated C2C12s exposed to 48-hour CM for 48 hours, normalized to GAPDH. Error bars represent standard deviation of the mean (**D**). Quantification of GFP intensity in HEK-PGFP reporter cells exposed to 48-hour CM, quantified at 0 and 24 hours of CM exposure. Adjusted p-values represent Holm-Bonferroni adjusted p-values of paired t-tests, n = 8 per group (**E**).

### BC-CM induces mitochondrial dysfunction and defects in lipid metabolism

We next asked whether BC-CM induced functional alterations in differentiated muscle cells that could be reflective of the increased muscle fatigability seen in women with BC and BC animal models. We found that BC-CM induced substantial changes in aerobic metabolism (**Figure 2A**), specifically leading to a reduction of oxygen consumed in ATP generation (**Figure 2B**) with inconsistent or negative results in other aerobic parameters across 5 independent experiments (not shown). CM from the luminal EO771 BC cell line repressed aerobic metabolism without altering glycolytic function, while the HER2-overexpressing NF639 line repressed both aerobic metabolism and glycolytic functions (**Figure 2C**). This reduction in aerobic metabolism does not appear to be related to mitochondrial biogenesis, as suggested by no alteration in mitochondrial DNA copy number in CM-treated myotubes (**Figure 2D**).

**Figure 2:**
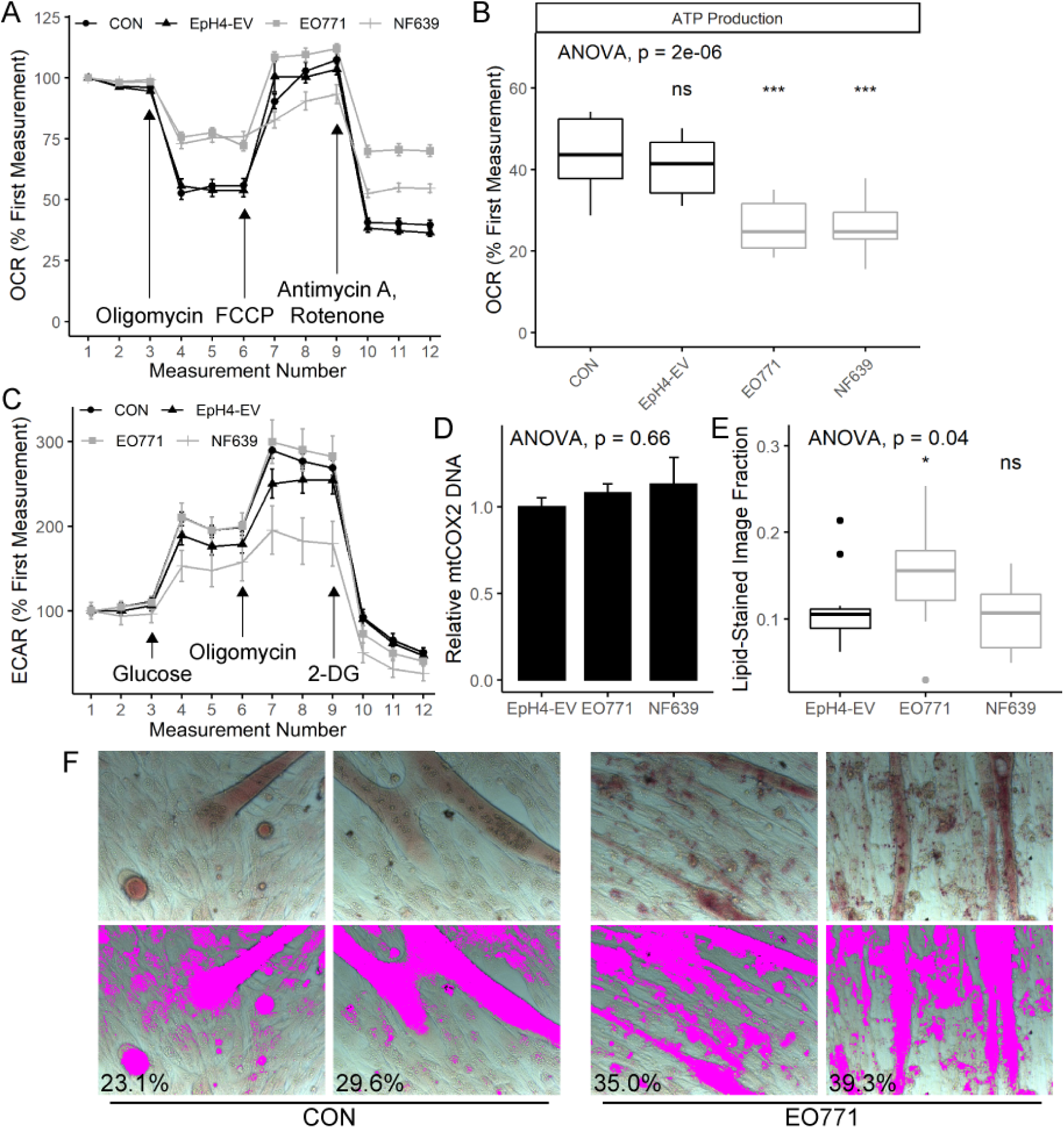
BC-CM induces mitochondrial dysfunction and defects in lipid metabolism. Normalized oxygen consumption rate (OCR) of 2 day differentiated C2C12 myotubes treated with 48-hour CM for 48 hours, quantified using the Seahorse XF Mito Stress Test. Error bars represent standard deviation of the mean, n = 10 per group (**A**). OCR associated with aerobic ATP production from A (**B**). Extra-cellular acidification rate of 2 day differentiated C2C12 myotubes treated with 48-hour CM for 48 hours, quantified using the Seahorse XF Glycolysis Stress Test. Error bars represent standard deviation of the mean, n = 12 per group, except n_NF639_ = 10 (**C**). Relative quantitation of mtCOX2 DNA in 5 day differentiated C2C12s exposed to 48-hour CM for 24 hours, normalized to 18S gDNA, n = 8 per group (**D**). Oil red o image quantification from 4 day differentiated C2C12s treated with 48-hour CM for 72 hours, represented as the fraction of each image stained red with oil red o, n = 8 per group (**E**). Representative images from an independent oil red o experiment conducted under identical conditions to E, 40X magnification (**F**).

Because lipotoxicity-induced mitochondrial dysfunction has been reported in various clinical contexts (22-25), and because individuals with cancer have been shown to have increased lipid content in muscle (26,27), we assessed intramyocellular lipid concentration in response to BC-CM using oil red o staining and automated image quantification. Using this methodology, we repeatedly observed a significant increase in intramyocellular lipid staining with EO771-CM, but in no experiment did we observe an increase in response to NF639-CM (**Figures 2E – 2F**).

### Exogenous PPARG expression rescues BC-CM-induced mitochondrial dysfunction and lipid accumulation

To determine whether or not the metabolic alterations induced in skeletal muscle by BC-derived factors are regulated by PPARG, we first attempted to generate stable myogenic cell lines expressing exogenous PPARG constructs or shRNA against PPARG using lentiviral infection and subsequent selection. Consistent with previous reports (28), genetic modification of PPARG in C2C12 myoblasts resulted in cell lines that rapidly lost differentiation competence, within five passages from lentiviral infection. Despite these limitations, early passage cells expressing exogenous PPARG (**Figure 3A**) were used to ascertain whether PPARG is involved in the response of myotubes to BC-derived factors. In these experiments, it was found that cells expressing exogenous PPARG were resistant to both BC-CM-induced repression of aerobic ATP generation (**Figure 3B**) as well as BC-CM-induced lipid accumulation (**Figures 3C – 3D**). Cells stably expressing shRNA against PPARG lost differentiation capacity immediately after lentiviral infection and could therefore not be used to determine if PPARG ablation phenocopies the effect of BC-CM in the context of differentiated muscle cells.

**Figure 3:**
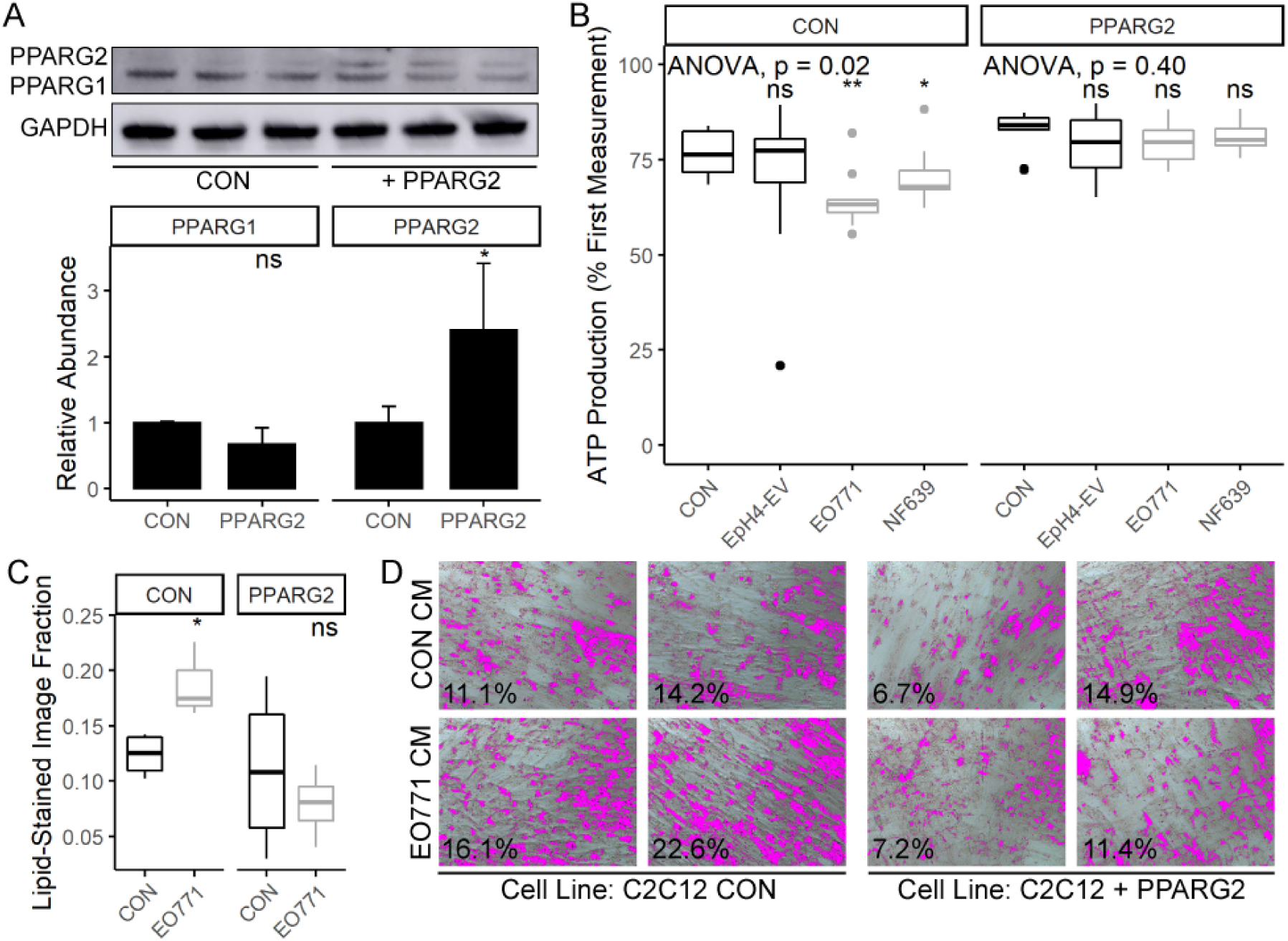
Exogenous PPARG expression rescues BC-CM-induced mitochondrial dysfunction and lipid accumulation. Western blotting analysis of PPARG expression in C2C12s infected with lentiviral particles carrying pLenti-C-PPARG2-mGFP-P2A-Puro (+PPARG2) or pLenti -mGFP-P2A-Puro (CON) after 5 days of differentiation (**A**). OCR associated with aerobic ATP production in 2 day differentiated CON or +PPARG2 cells exposed to 48-hour CM for 48 hours, quantified using the Seahorse XF Mito Stress Test, n = 10 per group (**B**). Oil red o image quantification from 4 day differentiated CON or +PPARG2 cells treated with 48-hour CM for 72 hours, represented as the fraction of each image stained red with oil red o, n = 6 per group (**C**). Representative images from the experiment quantified in C, 20X magnification (**D**).

### PPARG agonist rosiglitazone rescues BC-CM-induced lipid accumulation but fails to rescue BC-CM-induced mitochondrial dysfunction and PPAR repression

Finally, we pharmacologically targeted PPARG to overcome BC-induced mitochondrial dysfunction and lipid accumulation, using the potent PPARG agonist rosiglitazone (rosi) of the thiazolidinedione (TZD) drug class. Rosiglitazone did induce PPAR-mediated transcription in the absence of BC-CM (**Figure 4A**), but it was unable to rescue BC-CM’s repression of PPAR-mediated transcription (**Figure 4B**). Unsurprisingly then, this agonist did not rescue BC-induced mitochondrial dysfunction (**Figure 4C**). However, rosiglitazone did potently repress BC-induced lipid accumulation (**Figure 4D**). These data indicate that the mechanism by which rosiglitazone prevents BC-induced lipid accumulation is independent of PPARG’s transcriptional activity, and that BC-induced lipid accumulation is not the cause of BC-induced mitochondrial dysfunction in this model system.

**Figure 4:**
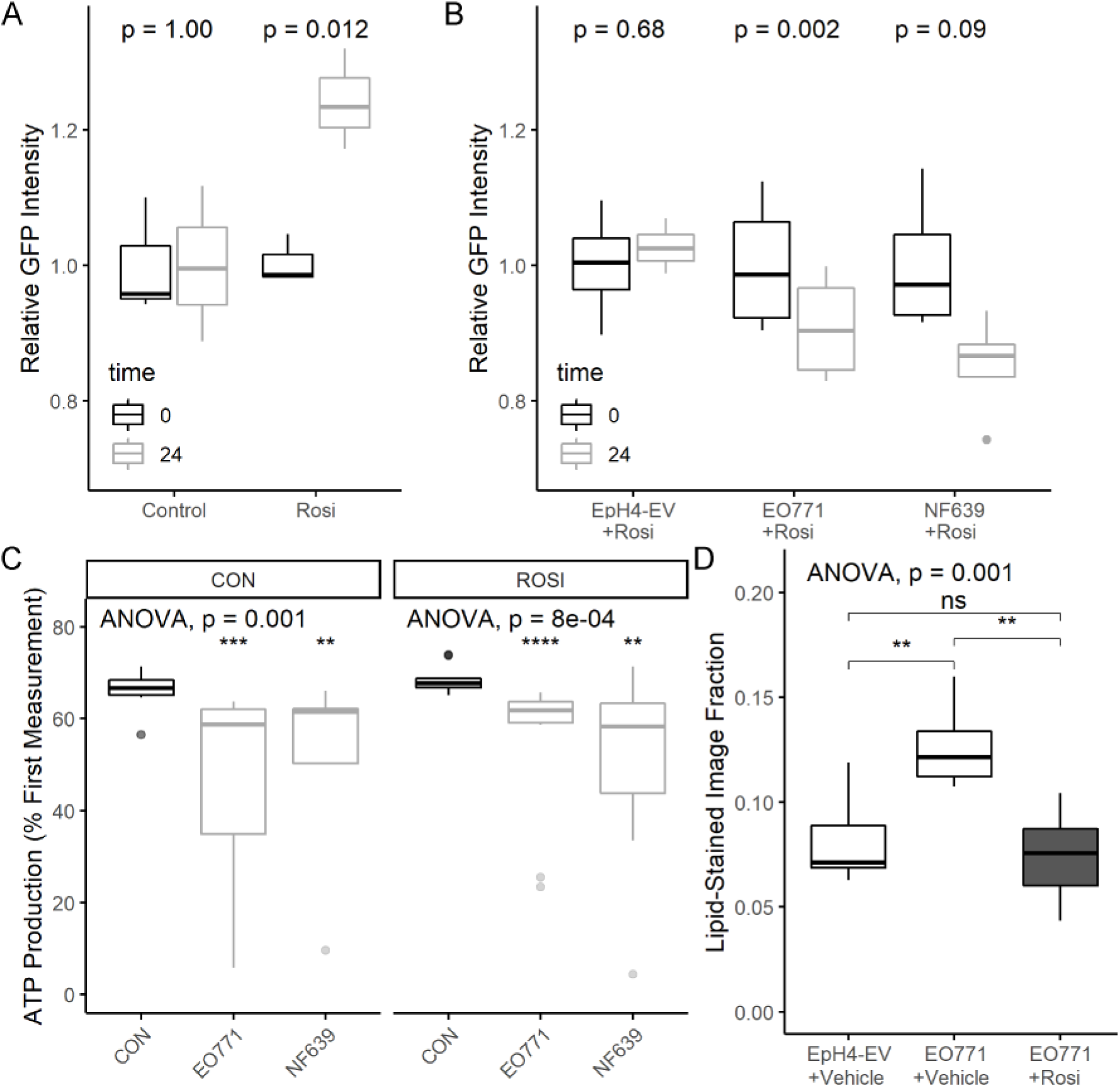
PPARG agonist rosiglitazone fails to rescue BC-CM-induced mitochondrial dysfunction and PPAR repression, but rescues BC-CM-induced lipid accumulation. Quantification of GFP intensity in HEK-PGFP reporter cells exposed to 10 uM rosiglitazone (rosi) or DMSO (control), quantified at 0 and 24 hours of drug exposure, n = 3 per group (**A**). Quantification of GFP intensity in HEK-PGFP reporter cells exposed to 48-hour CM in the presence of 10 µM rosiglitazone, quantified at 0 and 24 hours of CM exposure. Adjusted p-values represent Holm-Bonferroni adjusted p-values of paired t-tests, n = 4 per group (**B**). OCR associated with aerobic ATP production in 2 day differentiated C2C12s cells exposed to 48-hour CM +/- rosiglitazone for 48 hours, quantified using the Seahorse XF Mito Stress Test, n = 10 per group, except n_NF639+rosi_ = 7 (**C**). Oil red o image quantification from 4 day differentiated C2C12 cells treated with 48-hour CM for 72 hours, represented as the fraction of each image stained red with oil red o, n = 8 per group (**D**).

## DISCUSSION

In the present study, we have identified that BC-derived factors can significantly alter metabolic function and PPAR activity in skeletal muscle cells without the involvement of immune or stromal cell mediators, indicating that BC cells secrete a factor that induces these effects. Numerous publications have characterized changes in serum contents of women with BC, including changes in miRNAs, lipids, and proteins (29-33), though the exact sources of many of these factors remain unknown. To our knowledge, this report is the first showing that BC-CM induces alterations in skeletal muscle metabolic function, with previous studies focusing on BC-CM’s effect on immune cells, fibroblasts, endothelial cells, or normal mammary epithelium (34-39), or on transcriptional changes induced in skeletal muscle (40).

It is well established that BC cells can regulate both local and systemic immune functions (41,42), and that inflammatory signaling is intrinsically linked to cancer-associated skeletal muscle wasting (43,44). Thus, a reasonable assumption is that BC’s influence on immune cells is the cause of BC-induced skeletal muscle fatigue. This study contradicts this assumption by showing that no immunological mediators are required for BC cells to induce significant alterations in skeletal muscle metabolic function. This study also contradicts recent reports of negative results in other cancer types, where CM from cancer cells was unable to directly induce mitochondrial deficits in differentiated skeletal muscle (45), indicating that different cancer types induce skeletal muscle dysfunction via different mechanisms. While we cannot conclusively state that mitochondrial dysfunction or lipotoxicity are the root causes of BC-induced skeletal muscle dysfunction, it is reasonable to hypothesize a causal link that should be further investigated. Most likely, there are numerous mechanisms contributing to skeletal muscle dysfunction that would need to be simultaneously targeted to provide symptom control.

The mechanism by which BC cells induce metabolic dysfunction in this model system appears to be related to signaling by the PPAR proteins, as evidenced by the rescue of mitochondrial dysfunction and lipid content provided by exogenous expression of PPARG. This finding provides support for the translational relevance of this *in vitro* model system, as PPARG has been previously identified as a potential key regulator of BC-induced skeletal muscle fatigue using both human data and data generated in the patient-derived orthotopic xenograft model of BC (20,46). Additionally, this finding illuminates potential therapeutic modalities using the numerous FDA-approved agents that target the PPAR proteins, including drugs with various specificities for the three PPAR isoforms. In support of this possibility, we show here that the PPARG-specific agonist rosiglitazone completely reversed BC-CM’s effect on skeletal muscle lipid accumulation. Interestingly, this result does not appear to be due to rosiglitazone activating PPAR-mediated transcription, suggesting that the effect is mediated by non-canonical functions of PPARG.

While we were unable to rescue BC-induced mitochondrial dysfunction using rosiglitazone, it is possible that pharmacological agents targeting PPAR-alpha, PPAR-delta, or a combination of the three isoforms could overcome these effects of BC-derived factors. Alternatively, it is possible that the mitochondrial dysfunction induced by BC-CM is due to alteration of ligand-independent functions of PPARG, which would be overcome by the addition of PPARG protein but not by exogenous ligand. Numerous reports confirm the ability of PPARG to function as a transcription factor in the absence of ligand binding (47-50), and there is evidence of PPARG having ligand-independent functions that are also independent of its transactivation domain (51). Pharmacological strategies to target these unique functions could include non-TZD insulin-sensitizing agents or epigenetic modifiers.

Mechanistically, muscle fatigue can result from a myriad of factors, but the underlying ability of the contractile elements to maintain force production relies upon the muscle’s ability to resynthesize ATP following stimulus. We have observed dysregulation of metabolically significant PPAR transcriptional networks in the skeletal muscle of BC-PDOX mice (20), as well as a significant reduction in ATP from mitochondria isolated from skeletal muscle of these mice^1^ [unpublished observations]. In support of these murine observations, proteomics analysis in the skeletal muscle of HER2^+^ BC patients revealed lower protein expression of nearly every component of the mitochondrial electron transport chain (20) and a correspondingly lower ATP content in muscle biopsies from BC-patients of multiple tumor subtypes^2^ [unpublished observations]. These data clearly demonstrate mitochondrial dysfunction in muscle of patients with BC, which will directly impact the fatigability of skeletal muscle. The results presented herein expand on our previous studies and suggest this metabolic dysfunction is a direct result of BC-derived factors. Intriguingly, we demonstrate that this mitochondrial dysfunction is accompanied by increased intracellular lipid deposition in myotubes exposed to BC-CM. Intramuscular lipid deposition has also been observed in type 2 diabetes (52) and smoking-induced insulin resistance (53). As the downregulation of PPARG expression and transcriptional activity is associated with the development of insulin resistance (54) and many PPAR transcriptional targets are mitochondrial lipid transport proteins (21), we propose that a decreased capacity for mitochondrial lipid import may contribute to the observed phenotype of BC-induced lipid accumulation, which may then contribute to insulin resistance. Finally, aberrant intramyocellular lipid deposition can in many cases contribute to lipotoxicity characterized by the accumulation of toxic lipid intermediates such as diacylglycerols and ceramides (55), increased production of ROS (56), and is associated with changes in autophagic dynamics (57,58), all of which are implicated in reduced mitochondrial function within skeletal muscle. The sum of these observations suggest that skeletal muscle lipotoxicity in breast cancer could play a major role in the phenotype of BC induced skeletal muscle fatigue independent of mitochondrial dysfunction, and future studies should investigate this potential link in greater detail.

In summary, our results confirm the ability of BC-derived factors to directly alter metabolic function in skeletal muscle cells without the involvement of other cell types. These BC-CM-induced phenotypes in skeletal muscle could be enacted by a variety of secreted factors, including miRNAs, exosomes, or proteins, and appears to be mediated via signaling of the PPAR proteins, though perhaps not through their canonical functions as ligand-activated transcription factors. Specifically, BC-derived factors have the ability to repress the transcriptional activity of PPARG, and this effect is not due to downregulation of PPARG protein abundance. Additionally, repression of PPARG transcriptional activity induces mitochondrial dysfunction in muscle cells and a deficiency in ATP supply while also being associated with greater lipid accumulation. While PPARG protein levels are not affected in our model, basal protein expression of PPARG is relatively low in skeletal muscle, providing an explanation for the ability of exogenous PPARG to rescue these phenotypes while PPARG-agonists do not (**Figure 5**). These results provide support for previous publications implicating the PPAR proteins in BC-induced skeletal muscle fatigue and provide rationale for investigating PPAR agonists to improve quality of life in survivors of BC (20,46).

**Figure 5:**
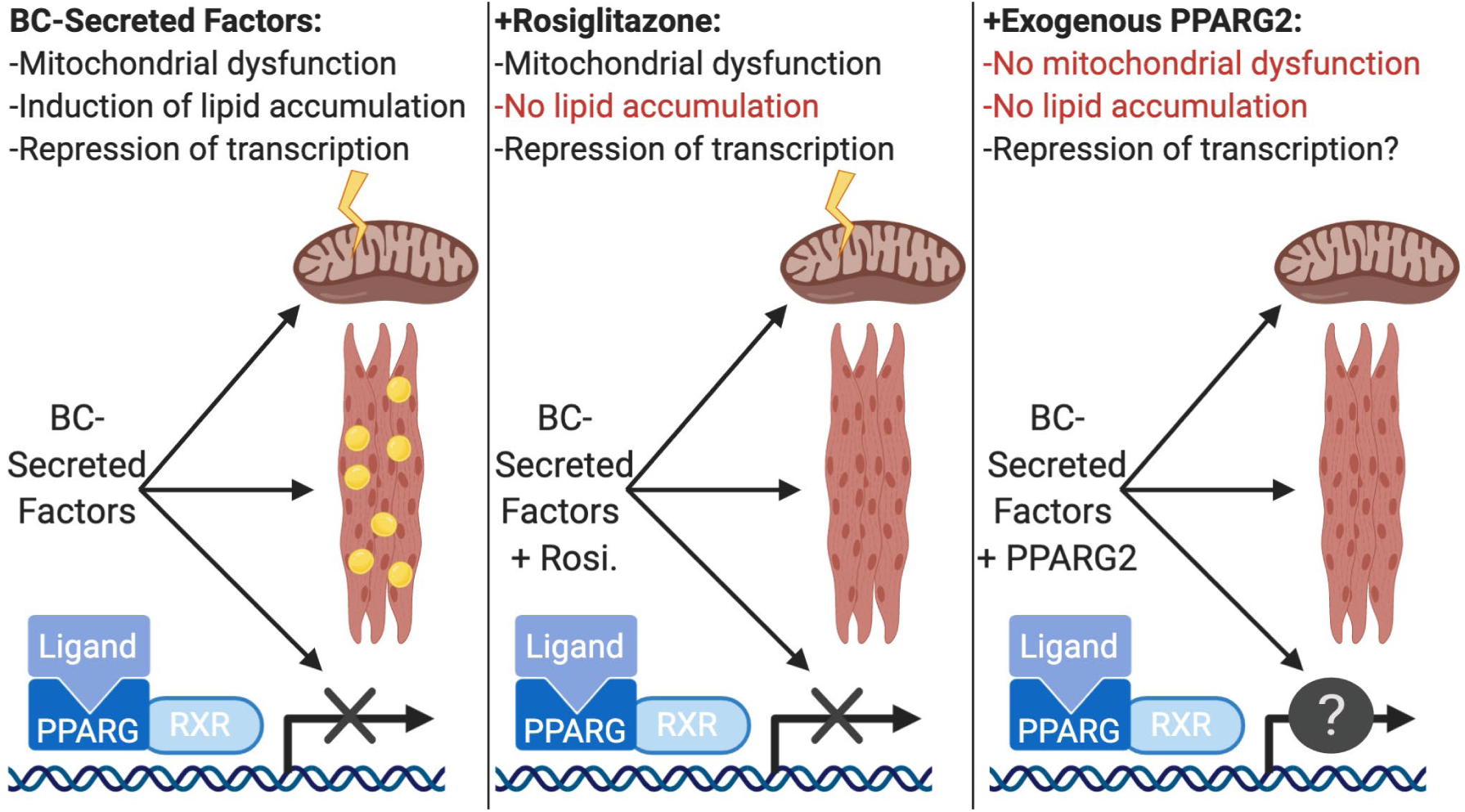
Summary of Findings. BC-derived factors induce several physiological changes in target cells *in vitro*, including mitochondrial dysfunction, lipid accumulation, and repression of PPAR-mediated transcription (**left**). The PPARG agonist Rosiglitazone reverses BC-induced lipid accumulation but does not reverse mitochondrial dysfunction or the observed repression of PPAR-mediated transcription (**center**). Expression of exogenous PPARG2 in myotubes reverses both BC-induced mitochondrial dysfunction and lipid accumulation. Figure created using BioRender (**right**).

## EXPERIMENTAL PROCEDURES

### Cell culture

Cell lines utilized include EpH4-EV (immortalized normal murine mammary epithelium), EO771 (murine luminal BC), NF639 (murine HER2/neu-overexpressing BC), HEK293 (human embryonic kidney), and C2C12 (murine myoblasts). All cell lines were obtained from ATCC (Virginia, USA), with the exception of EO771, which were obtained from Dr. Metheny-Barlow at Wake Forest University. All cell lines were cultured in DMEM (Thermo Fisher, Massachusetts, USA) supplemented with 10% heat inactivated fetal bovine serum (Atlanta Biologicals, Georgia, USA) and penicillin/streptomycin (Thermo Fisher) at 37°C with 6% CO_2_.

### Exogenous PPARG myogenic cell line

Lentiviral particles containing pLenti-C-PPARG2-mGFP-P2A-Puro or pLenti -mGFP-P2A-Puro were purchased from Origene (Maryland, USA). Titers were provided by the manufacturer. C2C12 cells were plated at 125,000 cells per well in 24-well plates and infected with a multiplicity of infection of 75 transforming units per cell with 8 µg · mL^-1^ polybrene in antibiotic-free DMEM. Media was changed 20 hours after infection. Starting 48 hours after infection, cells were cultured in 2 µg · mL^-1^ puromycin for 10 days, after which cells were maintained in 0.5 µg · mL^-1^ puromycin indefinitely. Uninfected control cells exhibited 100% death within 3 days of puromycin selection. Second passage cells were used in Western blotting analysis to assess expression of PPARG2 protein. All experiments with these cell lines were conducted within 5 passages of lentiviral infection as higher passage cells lost differentiation competence.

### Conditioned media (CM) collection

CM donor cells were plated at approximately 15% confluence in separate 10cm dishes for 48 hours. The 48-hour CM was then removed from all cell lines using a serological pipette, centrifuged at 1,500 RPM for 10 minutes, and the supernatants were collected via decanting into new centrifuge tubes. The collected CM was then diluted 1:3 in fresh growth media prior to application to recipient cells. CM was always applied to recipient cells within 2 hours of collection, in most cases within 15 minutes of collection **(Figure 1A)**. In some experiments, rosiglitazone (Millipore-Sigma, Massachusetts, USA) in dimethyl sulfoxide (DMSO) was added to the diluted conditioned media to final concentrations of 10 µM rosiglitazone and 0.1% DMSO prior to CM application to donor cells. In experiments where rosiglitazone was used, DMSO was added to control CM to 0.1% final concentration as a vehicle control.

### Western blotting

Differentiated C2C12s were lysed in 50 mM Tris-HCl (pH 6.8) with 2% sodium dodecyl sulfate and heated to 100°C for 3 cycles of 3 minutes each with vortexing and brief centrifugation between heating cycles. Lysates were diluted to a final concentration of 1 µg. µL^-1^ in NuPAGE LDS Sample Buffer (ThermoFisher Scientific) and 5% β-mercaptoethanol. 12 µg of total protein was loaded per well and resolved in NuPAGE Novex 4-12% Bis-Tris Gels (ThermoFisher Scientific). Proteins were transferred to nitrocellulose membrane, blocked for 1 hour in 1X tris-buffered saline (TBS), 0.1% Tween20, 5% milk followed by incubation with primary antibody overnight at 4°C in TBS-tween20 with 5% milk. Membranes were then washed 3 times in TBS + 0.1% Tween20 prior to application of appropriate secondary antibody (ThermoFisher Scientific) for 90 minutes at room temperature, and again prior to application of Pierce ECL Western Blotting Substrate (ThermoFisher Scientific). Relative band intensity was quantified using the GE Amersham Imager 600 (GE Healthcare Life Sciences, Marlborough, MA, USA) and normalized to GAPDH. Primary antibodies included PPARγ (#PA3-821A) and GAPDH (#2118S).

### CM metabolic analyses

C2C12 cells were plated into Agilent Seahorse XF96 (Agilent Technologies, California, USA) plates at 10,000 cells.well^-1^ and differentiated in 2% horse serum (Atlanta Biologicals) in DMEM with antibiotics for 48 hours. Meanwhile, EpH4-EV, EO771, NF639, and C2C12 cells were plated at approximately 15% confluence in separate 10cm dishes for 48 hours. 48-hour CM, prepared as described above, was then applied to the 2-day differentiated C2C12 cells in Seahorse assay plates for 48 hours (n=6-12 wells per treatment condition, noted in figure legends) prior to conducting the Agilent Seahorse XF Cell Mito Stress Test or Agilent Seahorse XF Glycolysis Stress Test protocol according to manufacturer’s instructions, using manufacturer’s recommended reagents.

### PPAR-reporter assays

HEK293 cells were transfected with PPRE-H2b-eGFP (59) (Addgene #84393) using Invitrogen Lipofectamine 3000 (Thermo Fisher), selected with 500 ng**·**uL^-1^ Gibco geneticin (Thermo Fisher) for 20 days, and flow-sorted to select the cells expressing GFP. The resulting HEK293-PPRE-H2b-eGFP cell line was plated at approximately 15% confluence in a 24-well plate. The following day, cells were imaged using the BioTek Cytation 5 Cell Imaging Multi-Mode Reader (Agilent Technologies) to collect baseline GFP intensity, and 72-hour conditioned media from HEK293, EpH4-EV, EO771, and NF639 cell lines was applied to the 24-well plate, using individual wells as biological replicates (n=6 wells per treatment condition). CM donor cells were plated on D_0_ as described above, and HEK-PGFP cells were plated at approximately 15% confluence on Day 1 (D_1_). On D_2_, baseline GFP intensity was collected using the BioTek Cytation 5 Cell Imaging Multi-Mode Reader (Agilent Technologies), and subsequently 48-hour conditioned media from HEK293, EpH4-EV, EO771, and NF639 cell lines was applied to the 24-well plate (n=3-6 wells per treatment condition, noted in figure legends). HEK-PGFP cells were then incubated in CM under normal culture conditions and GFP intensity was quantified 24 hours later using identical imaging settings as the baseline measurements. Mean GFP intensity per well was calculated using Gen5 Microplate Reader and Imager Software (Agilent Technologies), obtaining a single value per well that represented the mean GFP intensity of all cells in the imaging field for that well.

### qRT-PCR for PPAR target genes

C2C12 cells were plated at approximately 90% confluence in 24-well plates (150,000 cells/well) on D_0_ and differentiated in 2% horse serum (Atlanta Biologicals) in DMEM with antibiotics. On D_3_, differentiation media was refreshed on the C2C12s, and CM donor cells were plated as described under “CM collection.” On D_4_, media on the differentiating C2C12s was changed back to normal growth media (10% FBS in DMEM with antibiotics). On D_5_, CM was collected and applied to the differentiated C2C12 cells for 2 or 24 hours, with n=3-8 wells per condition, as noted in figure legends. Total RNA was isolated from CM- and/or drug-treated cells using Trizol (ThermoFischer Scientific, Waltham, MA, USA) and established methods (60). 1200 µg of cDNA was produced using Invitrogen SuperScript III First-Strand Synthesis System (ThermoFisher Scientific) according to manufacturer’s protocol, and relative expression of selected genes was analyzed using SYBR Green PCR Master Mix (ThermoFisher Scientific) with the Applied Biosystems QuantStudio 6 Flex (ThermoFisher Scientific), using 60 ng cDNA per reaction. Primer efficiencies were determined to be between 85% and 115% and relative mRNA expression was calculated using the Pfaffl method (61) normalized to 18s rRNA. Primer3 was used to design qRT-PCR primers **(Supplementary Table 1)** (62).

### qRT-PCR for mitochondrial DNA

C2C12 cells were plated at approximately 90% confluence in 24-well plates (100,000 cells.well-1) on D0 and differentiated in 2% horse serum in Gibco DMEM with antibiotics. Differentiation and CM application was conducted as described under “qRT-PCR for PPAR target genes,” with CM applied for 24 hours and n=8 wells per condition. Total DNA was isolated from CM-treated cells using the DNeasy Blood & Tissue Kit (Qiagen, Hilden, Germany) according to manufacturer’s instructions. 40-60 ng of DNA in 1uL eluting buffer was used for qRT-PCR using Applied Biosystemcs TaqMan Universal PCR Master Mix (ThermoFisher Scientific) with the Applied Biosystems QuantStudio 6 Flex (ThermoFisher Scientific). Primer-probe sets were purchased from ThermoFisher Scientific (Applied Biosystems Gene Expression Assays, COX2 Mm03294838_g1 and 18S Mm03928990_g1). Relative COX2 mtDNA was quantified using the ΔΔCT method normalized to 18S gDNA.

### Oil red O

C2C12 cells were plated at approximately 90% confluence in 24-well plates (100,000 cells.well^-1^) on D_0_ and differentiated in 2% horse serum in Gibco DMEM with antibiotics. On D_2_, differentiation media was refreshed on the C2C12s, and CM donor plates were set up as described under “CM collection.” On D_4_, CM was collected and applied to differentiated C2C12s, and CM donor cells were split to allow for CM to be collected once more after 48 hours incubation. On D_6_, CM was again collected and was used to refresh the CM on the recipient cells. On D_7_, cells were fixed in 4% formaldehyde for 30 minutes, washed 3 times in distilled water, and made permeable with 60% isopropanol for 10 minutes. Oil red o stock solution had been previously suspended in 100% isopropanol and filtered using a 0.2 µm bottle-top filter. Oil red o stock solution was diluted to 60% in distilled water to create the working solution. Working solution was applied to the fixed and permeable cells for 10 minutes. Cells were washed 3x with distilled water and stored at 4°C until imaging, typically within 24 hours of staining. Images were taken using a Zeiss Axiovert 40 CFL microscope with a Zeiss Axiocam 105 color camera, running ZEN 2.3 software. Effort was made to take images at the center of each well to produce even illumination across images. All images within each experiment were taken using identical imaging settings. Images were analyzed using the *countcolors* package version 0.9.1 (63) in R version 3.6.1 (64), quantifying the area of each image within a 50% radius from true red.

### Statistical analyses, reproducibility, and rigor

Data analyses were conducted in R version 3.6.1 (64) using *ggpubr* version 0.2.4 (65) for data visualization and most statistical tests. Unless otherwise stated, all statistical tests used were unpaired, two-tailed t-tests for two-group comparisons and one-way ANOVA for multi-group comparisons. When multiple comparisons were used within an experiment, Holm-Bonferroni correction was applied to adjust p-values. Graphs represent data from a single, independent experiment. All experiments have been repeated at least twice with similar results, with most experiments being repeated more than 3 times. For all box and whisker plots: the total height of the box represent the interquartile range, the thick center line represents the median value, and the whiskers extend to the most extreme data point that is not an outlier. Outlier values are represented by a single dot beyond the whiskers.

## DATA AVAILABILITY

The data described herein are available upon request. Requests should be directed to Dr. Emidio Pistilli.

## FUNDING AND ADDITIONAL INFORMATION

This research was supported by the following: National Institute of General Medical Sciences of the National Institutes of Health under Award Number P20GM121322 (Lockman), American Cancer Society Institutional Research Grant 09-061-04 (Pistilli), the WVCTSI U54GM104942 (Hodder). Authors would like to acknowledge the following WVU Core Facilities for contributing to this work: Flow Cytometry and Single Cell Core Facility (S10OD016165); Mitochondria Core of the WVU Stroke CoBRE (P20GM109098); Mitochondria, Metabolism and Bioenergetics group (R01 HL-128485; Hollander and the Community Foundation for the Ohio Valley Whipkey Trust). Imaging experiments were performed in the West Virginia University Microscope Imaging Facility which has been supported by the WVU Cancer Institute, the WVU HSC Office of Research and Graduate Education, and NIH grants P20RR016440, P30GM103488 and P20GM103434.

### Disclaimer

The content is solely the responsibility of the authors and does not necessarily represent the official views of the National Institutes of Health or other funding agencies.

## CONFLICTS OF INTEREST

The authors declare that they have no conflicts of interest with the contents of this article.

## ABBREVIATIONS

BC: Breast cancer
CM: conditioned media
PPAR: peroxisome-proliferator activated receptor
PPARG: peroxisome-proliferator activated receptor gamma
rosi: rosiglitazone
TBS: tris-buffered saline
TZD: thiazolidinedione

## TABLES

**Supplementary Table 1:**
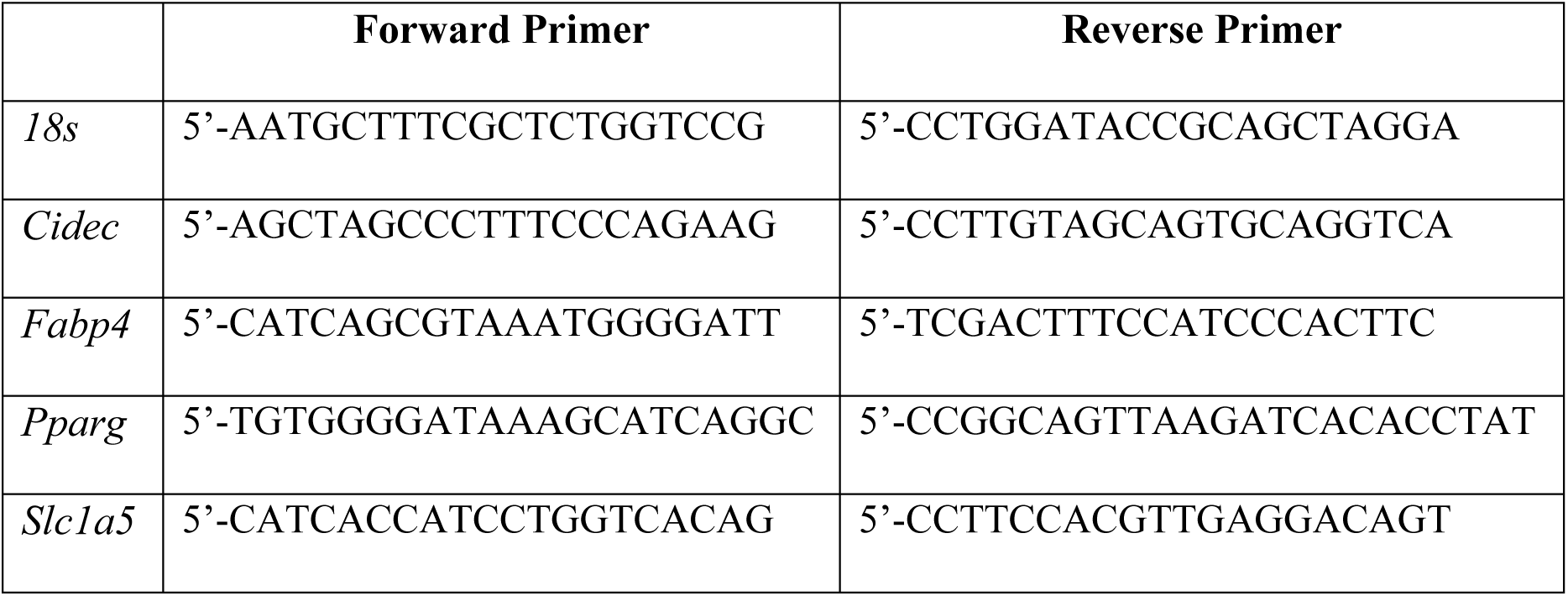
Primers used for qRT-PCR.

## FOOTNOTES

1 Wilson, H.E., Stanton D.A., Montgomery, C., Infante, A.M., Taylor, M., Hazard-Jenkins, H., Pugacheva, E.N., and Pistilli, E.E. (2019) Skeletal muscle reprogramming by breast cancer regardless of treatment history or tumor molecular subtype. *BioRxiv*, **doi:** https://doi.org/10.1101/810952.

2 Wilson, H.E., Stanton D.A., Montgomery, C., Infante, A.M., Taylor, M., Hazard-Jenkins, H., Pugacheva, E.N., and Pistilli, E.E. (2019) Skeletal muscle reprogramming by breast cancer regardless of treatment history or tumor molecular subtype. In Review, *npj Breast Cancer*, NPJBCANCER-00536-T.

## REFERENCES

1. Bower, J. E., Ganz, P. A., Desmond, K. A., Rowland, J. H., Meyerowitz, B. E., and Belin, T. R. (2000) Fatigue in breast cancer survivors: occurrence, correlates, and impact on quality of life. J Clin Oncol 18, 743–753

2. Cella, D., Lai, J. S., Chang, C. H., Peterman, A., and Slavin, M. (2002) Fatigue in cancer patients compared with fatigue in the general United States population. Cancer 94, 528–538

3. Curt, G. A., Breitbart, W., Cella, D., Groopman, J. E., Horning, S. J., Itri, L. M., Johnson, D. H., Miaskowski, C., Scherr, S. L., Portenoy, R. K., and Vogelzang, N. J. (2000) Impact of cancer-related fatigue on the lives of patients: new findings from the Fatigue Coalition. Oncologist 5, 353–360

4. Cella, D., Davis, K., Breitbart, W., Curt, G., and Coalition, F. (2001) Cancerrelated fatigue: prevalence of proposed diagnostic criteria in a United States sample of cancer survivors. J Clin Oncol 19, 3385–3391

5. Blesch, K. S., Paice, J. A., Wickham, R., Harte, N., Schnoor, D. K., Purl, S., Rehwalt, M., Kopp, P. L., Manson, S., and Coveny, S. B. (1991) Correlates of fatigue in people with breast or lung cancer. Oncol Nurs Forum 18, 81–87

6. Arndt, V., Stegmaier, C., Ziegler, H., and Brenner, H. (2006) A population-based study of the impact of specific symptoms on quality of life in women with breast cancer 1 year after diagnosis. Cancer 107, 2496–2503

7. Peters, K. B., West, M. J., Hornsby, W. E., Waner, E., Coan, A. D., McSherry, F., Herndon, J. E., 2nd, Friedman, H. S., Desjardins, A., and Jones, L. W. (2014) Impact of health-related quality of life and fatigue on survival of recurrent highgrade glioma patients. J Neurooncol 120, 499–506

8. Wang, X. S., and Woodruff, J. F. (2015) Cancer-related and treatment-related fatigue. Gynecol Oncol 136, 446–452

9. Groenvold, M., Petersen, M. A., Idler, E., Bjorner, J. B., Fayers, P. M., and Mouridsen, H. T. (2007) Psychological distress and fatigue predicted recurrence and survival in primary breast cancer patients. Breast Cancer Res Treat 105, 209–219

10. Prado, C. M., Baracos, V. E., McCargar, L. J., Reiman, T., Mourtzakis, M., Tonkin, K., Mackey, J. R., Koski, S., Pituskin, E., and Sawyer, M. B. (2009) Sarcopenia as a determinant of chemotherapy toxicity and time to tumor progression in metastatic breast cancer patients receiving capecitabine treatment. Clin Cancer Res 15, 2920–2926

11. Kersten, S., Desvergne, B., and Wahli, W. (2020) Roles of PPARs in health and disease. Nature 405, 421–424

12. Sabatino, L., Fucci, A., Pancione, M., and Colantuoni, V. (2012) PPARG Epigenetic Deregulation and Its Role in Colorectal Tumorigenesis. PPAR Res 2012

13. Aronoff, S., Rosenblatt, S., Braithwaite, S., Egan, J. W., Mathisen, A. L., and Schneider, R. L. (2000) Pioglitazone hydrochloride monotherapy improves glycemic control in the treatment of patients with type 2 diabetes: a 6-month randomized placebo-controlled dose-response study. The Pioglitazone 001 Study Group.

14. Chandra, S., and Pahan, K. (2019) Gemfibrozil, a Lipid-Lowering Drug, Lowers Amyloid Plaque Pathology and Enhances Memory in a Mouse Model of Alzheimer’s Disease via Peroxisome Proliferator-Activated Receptor α.

15. Lehrke, M., and Lazar, M. A. (2005) The many faces of PPARgamma. Cell 123, 993–999

16. Fan, W., and Evans, R. (2015) PPARs and ERRs: molecular mediators of mitochondrial metabolism. Curr Opin Cell Biol 33, 49–54

17. Auwerx, J., Cock, T. A., and Knouff, C. (2003) PPAR-gamma: a thrifty transcription factor. Nucl Recept Signal 1, e006

18. Torra, I. P., Chinetti, G., Duval, C., Fruchart, J. C., and Staels, B. (2001) Peroxisome proliferator-activated receptors: from transcriptional control to clinical practice. Curr Opin Lipidol 12, 245–254

19. Berger, J., and Moller, D. E. (2002) The mechanisms of action of PPARs. Annu Rev Med 53, 409–435

20. Wilson, H. E., Rhodes, K. K., Rodriguez, D., Chahal, I., Stanton, D. A., Bohlen, J., Davis, M., Infante, A. M., Hazard-Jenkins, H., Klinke, D. J., Pugacheva, E. N., and Pistilli, E. E. (2018) Human breast cancer xenograft model implicates peroxisome proliferator-activated receptor signaling as driver of cancer-induced muscle fatigue. Clinical Cancer Research, clincanres.1565.2018

21. Fang, L., Zhang, M., Li, Y., Liu, Y., Cui, Q., and Wang, N. (2016) PPARgene: A Database of Experimentally Verified and Computationally Predicted PPAR Target Genes. PPAR Res 2016, 6042162

22. Schrauwen, P., Schrauwen-Hinderling, V., Hoeks, J., and Hesselink, M. K. (2010) Mitochondrial dysfunction and lipotoxicity. Biochim Biophys Acta 1801, 266–271

23. Kumar, B., Kresge Eye Institute Wayne State University, D., Michigan, United States, Kowluru, A., Pharmaceutical Sciences, W. S. U., Detroit, Michigan, United States, β-Cell Biochemistry Laboratory, J. D. D. V. M. C., Detroit, Michigan, United States, Kowluru, R. A., and Kresge Eye Institute Wayne State University, D., Michigan, United States. (2020) Lipotoxicity Augments Glucotoxicity-Induced Mitochondrial Damage in the Development of Diabetic Retinopathy. Investigative Ophthalmology & Visual Science 56, 2985–2992

24. Yang, L., Wei, J., Sheng, F., and Li, P. (2020) Attenuation of Palmitic Acid–Induced Lipotoxicity by Chlorogenic Acid through Activation of SIRT1 in Hepatocytes - Yang - 2019 - Molecular Nutrition & Food Research - Wiley Online Library.

25. van de Weijer, T., Department of Human Biology, M. U. M. C., PO Box 616, 6200 MD Maastricht, The Netherlands, Schrauwen-Hinderling, V. B., Department of Radiology, S. o. N. a. M., NUTRIM, Maastricht University Medical Centre, The Netherlands, Schrauwen, P., and Department of Human Biology, M. U. M. C., PO Box 616, 6200 MD Maastricht, The Netherlands. (2020) Lipotoxicity in type 2 diabetic cardiomyopathy. Cardiovascular Research 92, 10–18

26. Stephens, N. A., Skipworth, R. J., Macdonald, A. J., Greig, C. A., Ross, J. A., and Fearon, K. C. (2011) Intramyocellular lipid droplets increase with progression of cachexia in cancer patients. J Cachexia Sarcopenia Muscle 2, 111–117

27. Gray, C., MacGillivray, T. J., Eeley, C., Stephens, N. A., Beggs, I., Fearon, K. C., and Greig, C. A. (2011) Magnetic resonance imaging with k-means clustering objectively measures whole muscle volume compartments in sarcopenia/cancer cachexia. Clin Nutr 30, 106–111

28. Singh, J., Verma, N. K., Kansagra, S. M., Kate, B. N., and Dey, C. S. (2007) Altered PPARgamma expression inhibits myogenic differentiation in C2C12 skeletal muscle cells. Mol Cell Biochem 294, 163–171

29. Hannafon, B. N., Trigoso, Y. D., Calloway, C. L., Zhao, Y. D., Lum, D. H., Welm, A. L., Zhao, Z. J., Blick, K. E., Dooley, W. C., and Ding, W. Q. (2016) Plasma exosome microRNAs are indicative of breast cancer. Breast Cancer Res 18, 90

30. Tozzoli, R., D’Aurizio, F., Falcomer, F., Basso, S. M., and Lumachi, F. (2016) Serum Tumor Markers in Stage I-II Breast Cancer. Med Chem 12, 285–289

31. Nunez, C. (2019) Blood-based protein biomarkers in breast cancer. Clin Chim Acta 490, 113–127

32. Delimaris, I., Faviou, E., Antonakos, G., Stathopoulou, E., Zachari, A., and Dionyssiou-Asteriou, A. (2007) Oxidized LDL, serum oxidizability and serum lipid levels in patients with breast or ovarian cancer. Clin Biochem 40, 1129–1134

33. Schwarzenbach, H. (2017) Clinical Relevance of Circulating, Cell-Free and Exosomal microRNAs in Plasma and Serum of Breast Cancer Patients. Oncol Res Treat 40, 423–429

34. Amirfakhri, S., Salimi, A., and Fernandez, N. (2015) Effects of Conditioned Medium from Breast Cancer Cells on Tlr2 Expression in Nb4 Cells. - PubMed - NCBI. Asian Pac J Cancer Prev 16, 8445–8450

35. Guo, J., Liu, C., Zhou, X., Xu, X., Deng, L., Li, X., and Guan, F. (2017) Conditioned Medium from Malignant Breast Cancer Cells Induces an EMT-Like Phenotype and an Altered N-Glycan Profile in Normal Epithelial MCF10A Cells. In Int J Mol Sci. pp

36. Wessels, D. J., Pradhan, N., Park, Y. N., Klepitsch, M. A., Lusche, D. F., Daniels, K. J., Conway, K. D., Voss, E. R., Hegde, S. V., Conway, T. P., and Soll, D. R. (2019) Reciprocal signaling and direct physical interactions between fibroblasts and breast cancer cells in a 3D environment. PLoS One 14, e0218854

37. Sousa, S., Brion, R., Lintunen, M., Kronqvist, P., Sandholm, J., Monkkonen, J., Kellokumpu-Lehtinen, P. L., Lauttia, S., Tynninen, O., Joensuu, H., Heymann, D., and Maatta, J. A. (2015) Human breast cancer cells educate macrophages toward the M2 activation status. Breast Cancer Res 17, 101

38. Luengo-Gil, G., Gonzalez-Billalabeitia, E., Perez-Henarejos, S. A., Navarro Manzano, E., Chaves-Benito, A., Garcia-Martinez, E., Garcia-Garre, E., Vicente, V., and Ayala de la Pena, F. (2018) Angiogenic role of miR-20a in breast cancer. PLoS One 13, e0194638

39. Furlan, A., Vercamer, C., Heliot, L., Wernert, N., Desbiens, X., and Pourtier, A. (2019) Ets-1 drives breast cancer cell angiogenic potential and interactions between breast cancer and endothelial cells. Int J Oncol 54, 29–40

40. Chen, D., Goswami, C. P., Burnett, R. M., Anjanappa, M., Bhat-Nakshatri, P., Muller, W., and Nakshatri, H. (2014) Cancer affects microRNA expression, release, and function in cardiac and skeletal muscle. Cancer Res 74, 4270–4281

41. Bates, J. P., Derakhshandeh, R., Jones, L., and Webb, T. J. (2018) Mechanisms of immune evasion in breast cancer. In BMC Cancer. pp

42. Gatti-Mays, M. E., Balko, J. M., Gameiro, S. R., Bear, H. D., Prabhakaran, S., Fukui, J., Disis, M. L., Nanda, R., Gulley, J. L., Kalinsky, K., Sater, H. A., Sparano, J. A., Cescon, D., Page, D. B., McArthur, H., Adams, S., and Mittendorf, E. A. (2019) If we build it they will come: targeting the immune response to breast cancer. npj Breast Cancer 5, 1–13

43. Fearon, K., Strasser, F., Anker, S. D., Bosaeus, I., Bruera, E., Fainsinger, R. L., Jatoi, A., Loprinzi, C., MacDonald, N., Mantovani, G., Davis, M., Muscaritoli, M., Ottery, F., Radbruch, L., Ravasco, P., Walsh, D., Wilcock, A., Kaasa, S., and Baracos, V. E. (2011) Definition and classification of cancer cachexia: an international consensus. Lancet Oncol 12, 489–495

44. Fearon, K. C., From the Royal Infirmary of Edinburgh, E., United Kingdom (KCF), and the Ross Products Division, Abbott Laboratories, Columbus, OH (ACV and DSH), Voss, A. C., From the Royal Infirmary of Edinburgh, E., United Kingdom (KCF), and the Ross Products Division, Abbott Laboratories, Columbus, OH (ACV and DSH), Hustead, D. S., and From the Royal Infirmary of Edinburgh, E., United Kingdom (KCF), and the Ross Products Division, Abbott Laboratories, Columbus, OH (ACV and DSH). (2006) Definition of cancer cachexia: effect of weight loss, reduced food intake, and systemic inflammation on functional status and prognosis. The American Journal of Clinical Nutrition 83, 1345–1350

45. Guigni, B. A., van der Velden, J., Kinsey, C. M., Carson, J. A., and Toth, M. J. (2020) Effects of conditioned media from murine lung cancer cells and human tumor cells on cultured myotubes. Am J Physiol Endocrinol Metab 318, E22–e32

46. Bohlen, J., McLaughlin, S. L., Hazard-Jenkins, H., Infante, A. M., Montgomery, C., Davis, M., and Pistilli, E. E. (2018) Dysregulation of metabolic-associated pathways in muscle of breast cancer patients: preclinical evaluation of interleukin-15 targeting fatigue. J Cachexia Sarcopenia Muscle

47. Werman, A., Hollenberg, A., Solanes, G., Bjorbaek, C., Vidal-Puig, A. J., and Flier, J. S. (1997) Ligand-independent activation domain in the N terminus of peroxisome proliferator-activated receptor gamma (PPARgamma). Differential activity of PPARgamma1 and −2 isoforms and influence of insulin. J Biol Chem 272, 20230–20235

48. Jiang, X., Ye, X., Guo, W., Lu, H., and Gao, Z. (2014) Inhibition of HDAC3 promotes ligand-independent PPARgamma activation by protein acetylation. J Mol Endocrinol 53, 191–200

49. Hurtado, O., Ballesteros, I., Cuartero, M. I., Moraga, A., Pradillo, J. M., Ramirez-Franco, J., Bartolome-Martin, D., Pascual, D., Torres, M., Sanchez-Prieto, J., Salom, J. B., Lizasoain, I., and Moro, M. A. (2012) Daidzein has neuroprotective effects through ligand-binding-independent PPARgamma activation. Neurochem Int 61, 119–127

50. Al-Rasheed, N. M., Chana, R. S., Baines, R. J., Willars, G. B., and Brunskill, N. J. (2004) Ligand-independent activation of peroxisome proliferator-activated receptor-gamma by insulin and C-peptide in kidney proximal tubular cells: dependent on phosphatidylinositol 3-kinase activity. J Biol Chem 279, 49747–49754

51. Daniel, B., Nagy, G., Czimmerer, Z., Horvath, A., Hammers, D. W., Cuaranta-Monroy, I., Poliska, S., Tzerpos, P., Kolostyak, Z., Hays, T. T., Patsalos, A., Houtman, R., Sauer, S., Francois-Deleuze, J., Rastinejad, F., Balint, B. L., Sweeney, H. L., and Nagy, L. (2018) The Nuclear Receptor PPARgamma Controls Progressive Macrophage Polarization as a Ligand-Insensitive Epigenomic Ratchet of Transcriptional Memory. Immunity 49, 615-626.e616

52. Brons, C., and Grunnet, L. G. (2017) MECHANISMS IN ENDOCRINOLOGY: Skeletal muscle lipotoxicity in insulin resistance and type 2 diabetes: a causal mechanism or an innocent bystander? Eur J Endocrinol 176, R67–r78

53. Bergman, B. C., Perreault, L., Hunerdosse, D. M., Koehler, M. C., Samek, A. M., and Eckel, R. H. (2009) Intramuscular Lipid Metabolism in the Insulin Resistance of Smoking. In Diabetes. pp 2220–2227

54. Hevener, A. L., He, W., Barak, Y., Le, J., Bandyopadhyay, G., Olson, P., Wilkes, J., Evans, R. M., and Olefsky, J. (2003) Muscle-specific Pparg deletion causes insulin resistance. Nat Med 9, 1491–1497

55. Chavez, J. A., and Summers, S. A. (2003) Characterizing the effects of saturated fatty acids on insulin signaling and ceramide and diacylglycerol accumulation in 3T3-L1 adipocytes and C2C12 myotubes. Arch Biochem Biophys 419, 101–109

56. Rindler, P. M., Crewe, C. L., Fernandes, J., Kinter, M., and Szweda, L. I. (2013) Redox regulation of insulin sensitivity due to enhanced fatty acid utilization in the mitochondria. Am J Physiol Heart Circ Physiol 305, H634–643

57. Morales, P. E., Bucarey, J. L., and Espinosa, A. (2017) Muscle Lipid Metabolism: Role of Lipid Droplets and Perilipins. J Diabetes Res 2017, 1789395

58. Sarparanta, J., Garcia-Macia, M., and Singh, R. (2017) Autophagy and Mitochondria in Obesity and Type 2 Diabetes. Curr Diabetes Rev 13, 352–369

59. Degrelle, S. A., Shoaito, H., and Fournier, T. (2017) New Transcriptional Reporters to Quantify and Monitor PPARgamma Activity. PPAR Res 2017, 6139107

60. Pistilli, E. E., Jackson, J. R., and Alway, S. E. (2006) Death receptor-associated pro-apoptotic signaling in aged skeletal muscle. Apoptosis: an international journal on programmed cell death 11, 2115–2126

61. Pfaffl, M. W. (2001) A new mathematical model for relative quantification in real-time RT-PCR. Nucleic Acids Res 29, e45

62. Untergasser, A., Cutcutache, I., Koressaar, T., Ye, J., Faircloth, B. C., Remm, M., and Rozen, S. G. (2012) Primer3--new capabilities and interfaces. Nucleic Acids Res 40, e115

63. Weller, H. (2019) countcolors: Locates and Counts Pixels Within Color Range(s) in Images., R, https://CRAN.R-project.org/package=countcolors

64. (2019) R: A language and environment for statistical computing., R Foundation for Statistical C omputing

65. Kassambara, A. (2019) ggpubr: ‘ggplot2’ Based Publication Ready Plots. Comprehensive R Archive Network (CRAN)

